# Disorder and interfaces in proteins are two sides of the same coin

**DOI:** 10.1101/863571

**Authors:** Beatriz Seoane, Alessandra Carbone

## Abstract

The importance of unstructured biology has quickly grown during the last decades accompanying the explosion of the number of experimentally resolved structures. The idea that structural disorder might be a novel mechanism of protein interaction is widespread in the literature, although the number of statistically significant structural studies supporting this idea is surprisingly low. In this work, through a large-scale-analysis of all the crystallographic structures of the Protein Data Bank averaged over clusters of homologous sequences, we show clear evidences that both the (experimentally verified) interaction interfaces and the disordered regions are involving roughly the same amino-acids of the protein. And beyond, disordered regions appear to carry information about the location of alternative interfaces when the protein lies within complexes, thus playing an important role in determining the order of assembly of protein complexes.

---

The paradigm by which the protein function is determined by its three-dimensional structure is one of the basis of Molecular Biology. However, as long as the number of experimental structures increases, it gets clearer that many perfectly functional proteins either lack a well defined structure or they are largely unstructured^1–4^. These proteins are known as intrinsically disordered proteins (IDP) or regions (IDR), they are fairly abundant among known proteins^5^ (and ubiquitous in eukaryotic proteomes^6–8^) and pathologically present in severe illnesses^9^.

The increasing importance of the IDPs is calling for a reformulation of the structure-function paradigm itself^10, 11^, but the biological function of these IDPs is far from being understood. It is known that IDRs tend to enhance the protein flexibility^12^, which would increase the number of conformational structures and thus their promiscuity^13–15^. This agrees with the fact that IDRs are often involved in tasks associated to molecular recognition, including the gene regulation, folding assistance or cellular cycle control^4^. Nowadays it is believed that one of the main functions of these IDPs is precisely to facilitate the binding with other partners (other proteins, DNA, RNA or small molecules), so it is often invoked as a novel mechanism of protein interaction, even though the statistical evidence of this claim is rather scarce^16, 17^. In addition, IDRs often structure after the binding, suffering a so-called disorder-to-order transition^18, 19^, which makes them a hot target for drug discovery^20, 21^.

The vast majority of the large-scale computational studies exploring the biological function of IDRs rely on bioinformatics’ disorder predictors based on the amino-acid (AA) sequence to fill the gap between the amount of expected and observed disorder^22, 23^. This fact hampers the identification of universal mechanisms mainly for two reasons. First, because more than 60 different predictors, identifying alternative flavors of disorder (not necessarily mutually compatible), including meta-predictors, have been proposed^24, 25^. And second, because these predictive methods, in their majority, are developed using a reduced set of the total experimental information available about disorder (for instance, the information between the observed and missing regions in the X-ray crystallographic structures of the Protein Data Bank (PDB)^26^). This information often disagrees between the different structures of the same protein sequence (see Ref.^27^ and this manuscript), so a question of generality arises. Contrary to all those previous approaches, our conclusions rely exclusively on a direct analysis of all the crystallographic structures of the PDB.

In this work, we have explored the link between disorder regions (DRs) and interface regions (IRs) in a protein sequence by crossing structural data from homologous families of proteins and using all the structures available in the PDB. We observe that, whenever several structures for a given family are available, the union of all the DRs observed in them predicts relatively well the union of all the IRs, highlighting a direct and simple connection between both mechanisms. Furthermore, we show that the knowledge of the disorder of one particular structure gives information about the location of possible alternative interfaces and the order of assembly of complexes. We divide our presentation as follows. We begin with a description of our analysis, followed by the results based on the union of disorder. Finally, we discuss the role of the disorder at different stages of complex formation and show several examples where disorder predicts the assembly order.

## 1 Analysis procedure

The first step of our analysis is to compute all the IRs and DRs in all the X-ray crystallographic structures of the PDB. To build the IRs, we sum up all the protein-protein/DNA/RNA binding sites. For the DRs, we consider, not only the so-called missing residues, but also the residues with an anomalously high b-factor. The details of both definitions are discussed in Methods. This extended definition of disorder gathers, at the same time, the flexible and the amorphous parts of the chain. As a second step, we cluster together all the information from experiments of similar proteins (where a cluster will be labelled by the name of its representative chain structure in the form of PDB ID, underscore, chain name; see Methods). Finally, we compute the union of all the IRs and DRs (namely, UIRs and UDRs from now on) observed in the cluster by mapping all those regions in the representative’s sequence, and assigning to each of its AAs the label IR and/or DR whenever it was interface or disordered at least once in the cluster. The descriptions of the clustering and union procedures are given in Methods. We illustrate the pipeline of this procedure and an example of a cluster in Fig. 1.

**Figure 1:**
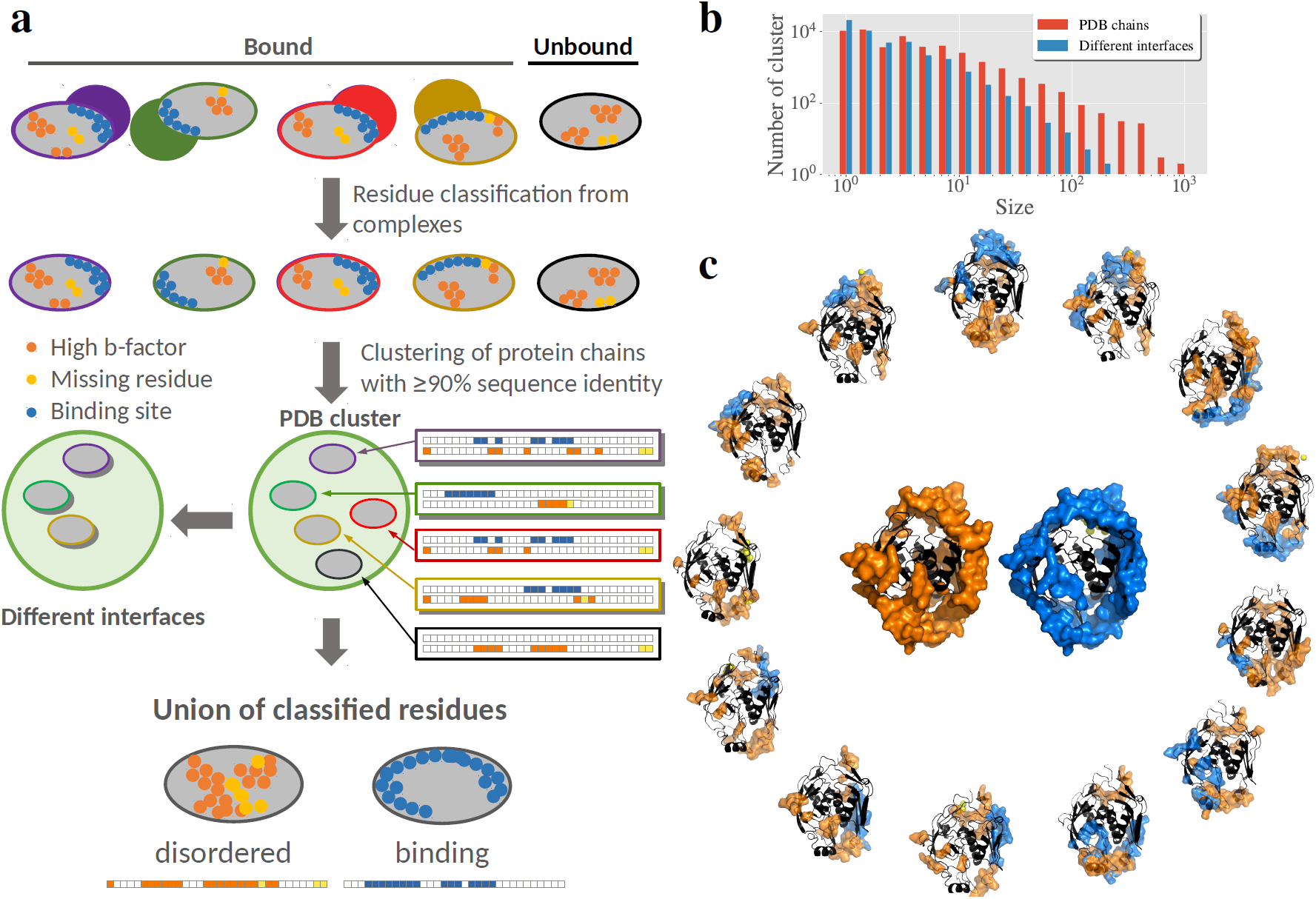
Pipeline of the analysis. **a.** Sketch of the analysis procedure: (i) We compute all binding (blue), missing (yellow) and high b-factor (orange) residues for all protein chains in the PDB. (ii) We cluster together chains with a common sequence identity ≥ 90%. (iii) For each cluster, we compute the UDR as the union of all AA flagged as high-b-factor or missing in at least one chain of the cluster, and same for the UIRs with the binding sites. Clusters contain a variable number of chains (5 here). Chains without any interface are considered as “unbound” (black structure here), and the rest as “bound”. Each cluster is characterised by the number of chains with different interfaces (3 here). **b.** Number of clusters with a particular size (number of PDB chain structures, red; and with a given NDI, blue). **c.** Analysis of cluster 6c9c A (PDB IDs for the structures are given in SI section S2). In the external wheel, we show one unbound structure and the chain structures containing the 12 different interfaces of this cluster: DRs in orange and IRs in blue. Missing residues are modelled as yellow spheres and localised close to those AA in the structure that are direct neighbours in the sequence. In the centre, UDR (orange) and UIR (blue) are mapped on the structure of the cluster’s representative.

In total, we have analysed *N*_c_ = 47102 clusters containing 354339 chains extracted from 134337 PDB protein structures. We say that a chain is *bound* if an interface can be measured with at least one other protein/DNA/RNA chain in the PDB complex, and *unbound* otherwise. 16605 of the total clusters contain one or more unbound chains, but only 6131 of them contain also bound structures. Our clusters are composed of a variable number of chains, having a large number of them (see Table S1 in the SI) just one. We found crucial for our analysis to identify each cluster by its number of different interfaces (NDI). We consider two interfaces as *different* if the number of non mutual AAs in the interfaces is larger than 5% of the sequence length. This means that two interfaces that are concentric or slightly displaced, but one significantly larger than the other, are counted as different interfaces. We show in Fig. 1–**b** the total number of clusters with a given size and NDI. We show a breakdown of the clusters composition and the full list of the clusters in Supplemental Information (SI) section S1.

## 2 Results

The DRs look qualitatively similar to the IRs in the example of Fig. 1-**c**, although an AA is not typically both disordered and belonging to an interface for the same structure (binding sites have generally a low b-factor, and by construction, they cannot be missing). This idea gets more quantitative when we compare the relative size of the UDRs, *r*_D_ = *N*_D_*/L* where *N*_D_ is the number of disordered sites and *L* the sequence length, to the relative size of the UIRs, *r*_I_ = *N*_I_*/L* with *N*_I_ the number of binding sites. We show the distribution of *r*_D_ and *r*_I_ among clusters of similar NDI (in logarithmic scale) in Fig. 2–**a**. The two kinds of regions follow very similar trends as NDI increases, a trend that is destroyed if we randomly displace the DRs measured at each experiment, because the UDR quickly covers the entire sequence (see Fig. S2, where prefix S means that it is found in the SI). This last observation confirms that the positions of the DRs are not random, but localised in well defined regions, just like the IRs.

**Figure 2:**
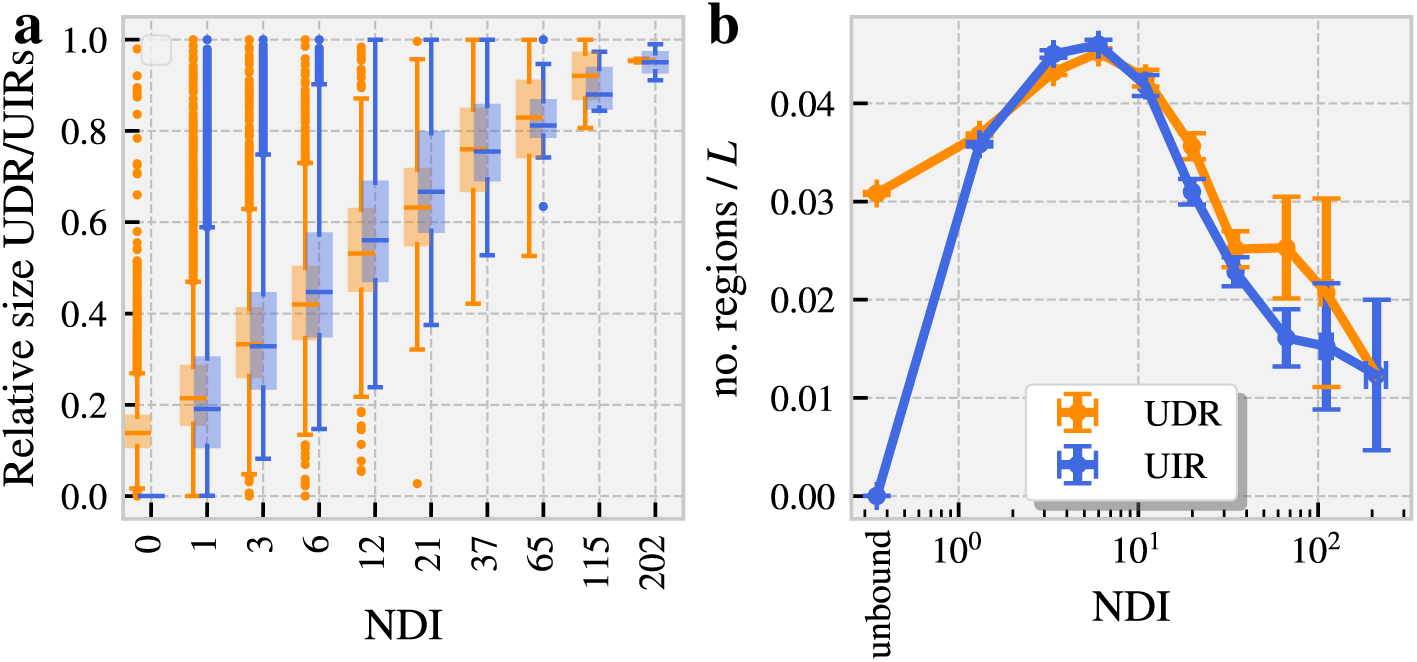
UDRs vs. UIRs using all the protein chains in the PDB. **a** Box-plot of the relative size of the UDR (orange) and UIR (blue) in the sequence, as function of the NDI using bins of equal size in logarithmic scale (in the x-label shows the bin’s central value). **b** Number of structurally connected regions in the UDR and UIR divided by the sequence length *L* and averaged over clusters of similar NDI (bins defined as in **a**). The first two points correspond to clusters composed uniquely by unbound structures. The error variables are the standard error of the mean (computed from the bins’ cluster-to-cluster fluctuations).

We can also compare the average number of distinct structurally connected regions in the UDRs and UIRs. To do so, we recursively group together all the AAs assigned as DR/IR located within a distance (in the structure) of ≤ 6Å to at least one of the other members of the group. As we show in Fig. 2–**b**, the number of DRs and IRs (normalised by *L*) follows, again, a surprisingly similar behaviour once averaged over all the clusters with a similar NDI: they both grow with the NDI up to approximately 6, moment at which regions start to superpose thus decreasing the number of regions. Quite interestingly, a random displacement of the groups of DRs in the UDR or a total randomisation of the disordered sites of the UDR along the sequence, lead different curves, as shown in Fig. S3.

### Disorder as predictor of interfaces

The UIRs and UDRs display a surprisingly similar behaviour as the clusters’ NDI grows, both in terms of relative sizes and number of structural regions. It is then natural to wonder whether both regions actually coincide at the same AAs after taking the union over all the cluster structures. In other words, if DRs can work as a predictor of the location of IRs. To test this idea, we need to count, cluster-by-cluster, the number of disordered residues that are also IR (true positives) and those that are not (false positives). And in the other way round, the number of residues that are neither DR nor IR (true negatives) and those that are IR but not DR (false negatives). We can use combinations of these 4 measures to test the quality of the predictor. We have considered 5 different metrics: (i) sensitivity (fraction of the total interface predicted), (ii) specificity (fraction of the non-interface region that is correctly predicted), (iii) accuracy (fraction of correct assignations among the total, both interface and not interface), (iv) precision or positive predictive value (fraction of the predicted sites that are also binding sites), and (v) F1-score (overall goodness of the prediction, an harmonic mean between sensitivity and precision). We show the definition of each estimator in Methods and, all the cluster values as function of their NDI in Fig. 3. The dots’ shade of grey reflects the density of cluster values in bins of similar NDI, and the black line, the value of the median within these bins. Clearly, the larger the NDI, the better the predictor, a result that one should expect considering that our data base of alternative interfaces is very incomplete.

**Figure 3:**
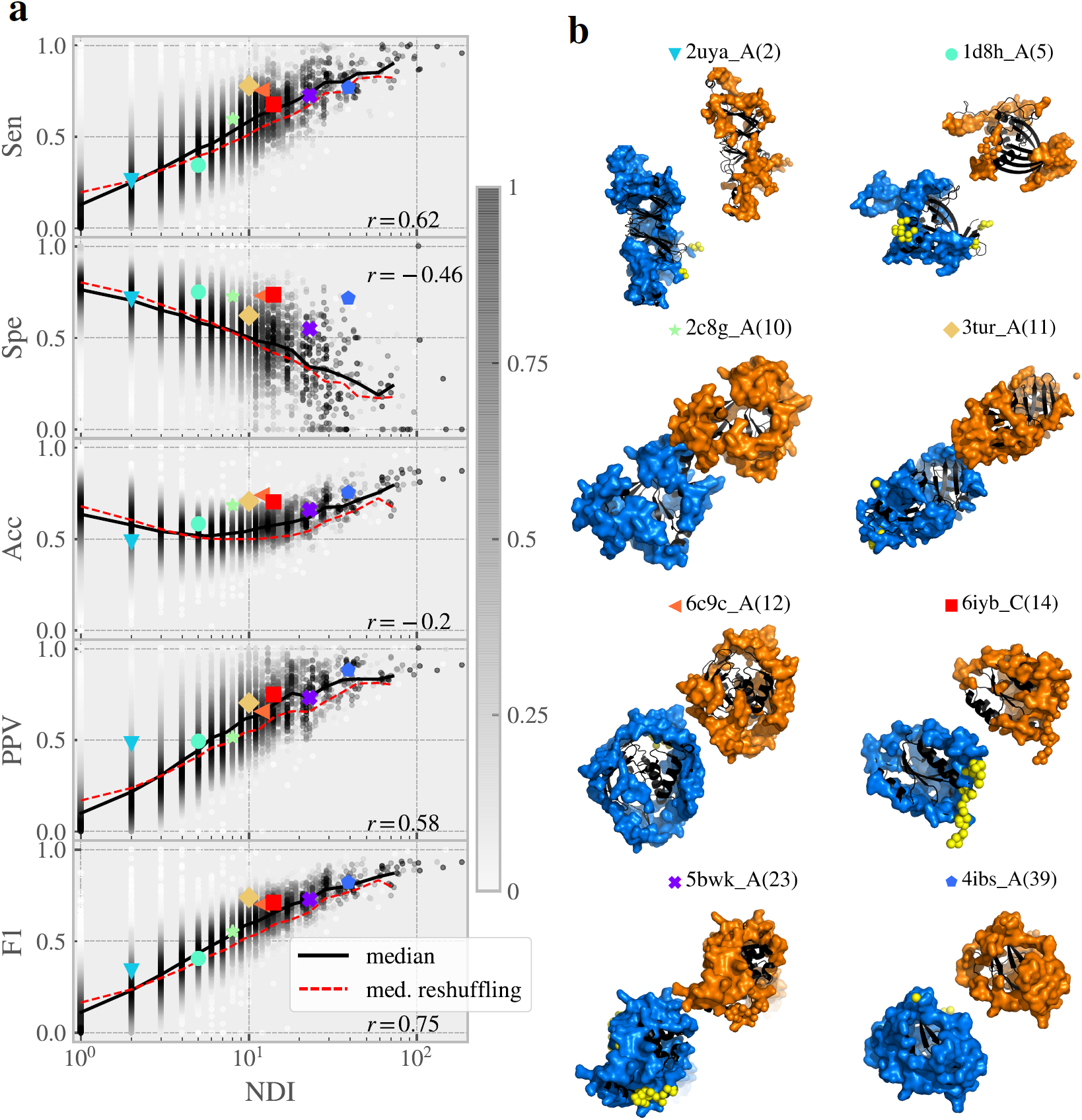
Disorder as interface predictor. **a** We test how reliable the prediction of interfaces based on the knowledge of the disorder is, for each of the clusters available in the PDB. We compute 5 different metrics of its performance [Sensitivity (Sen), Specificity (Spe), Accuracy (Acc), Precision (PPV), F1-score]. The measures for each cluster (dots) are plotted as function of its NDI. The white-to-black colour reflects the density of cluster points within bins of similar NDI (we considered 30 bins of equal width in log-scale). We also show, as a full-black-line, the median of all these measures computed in each bin (only for the bins that contain at least 10 clusters). With the sake of comparison, we include, as a red-dashed-line, the median among clusters of the expected estimators for a random reshuffling of the UDR. The correlation coefficient between each metric and the logarithm of the NDI for each cluster is shown on the bottom-right of each panel. **b** We show several snapshots of the protein structure of 8 clusters where the UIR and UDR in the cluster is shown as blue and orange surfaces, respectively. The PDB ID of the cluster representative and the cluster’s NDI is shown on the top, together with a big-colour point that allows us locate the cluster metrics evaluation in **a**.

Yet, since both the sizes of the DRs and IRs increase with the NDI, so does the goodness of a random-guess prediction of the same size than the real DR. For this reason, the estimator values are only meaningful if compared with those of a random reshuffling of the disordered sites along the sequence. At this point, let us stress that it is not a standard randomisation test. Indeed, a complete randomisation of the DRs extracted from the experiments leads to entirely disordered chains after the union, as shown in Fig. S2, which is not a reasonable prediction. At variance, we randomise only after having computed the union of the experimental DRs (which already contains some information about the number of interfaces in the cluster as we showed before). The expected metrics for the random case can be easily expressed, cluster by cluster, in terms of its *r*_D_ and *r*_I_, as we discuss in Methods. Other kinds of randomisation have been considered in SI section S6, giving very similar results. We also include the median of the random prediction in Fig. 3. We see that for all the estimators considered, the real disorder prediction is better than its randomised version as long as the NDI is larger than two, which is natural considering that DRs do not generally superpose with IRs in the same structures so such a prediction must be worse than random. The improvement over a random might look too moderate, but we show below that this is mostly related to the lack of information in our database, that is, because the unbound structures are missing and/or many of the predicted interfaces were not measured yet.

### Disorder and protein assembly

In the following, we will show evidences that DRs predict the possible interface regions allowed at the next step of complex assembly. That is, the DRs observed in the unbound forms of a protein give us information about the interaction sites of this protein with other partners, and the DRs observed in a complex, about the interfaces of this complexes with further partners. We illustrate this idea in the sketch of Fig. 4–**a**. The simplest statistical test of this hypothesis requires splitting our whole set of clusters according to the existence (or not existence) of unbound structures for that protein, so we can check the first step of assembly. With this aim, we compare, our 5 prediction metrics obtained using: (i) all the *N*_c_ clusters (as in Fig. 3), (ii) only the clusters with one or more unbound structures, (iii) clusters without any unbound structure, and finally, (iv) using only the DRs observed in the unbound structures to compute the UDR and all the structures for the UIR. The sizes of the UDRs vary with the group considered (specially for the predictions from the unbound), which affects the values of the metrics. To avoid this effect, we compare the performance of the disorder predictors via the relative improvement of the estimators with respect to random guess predictions. That is, cluster by cluster, we compute:

**Figure 4:**
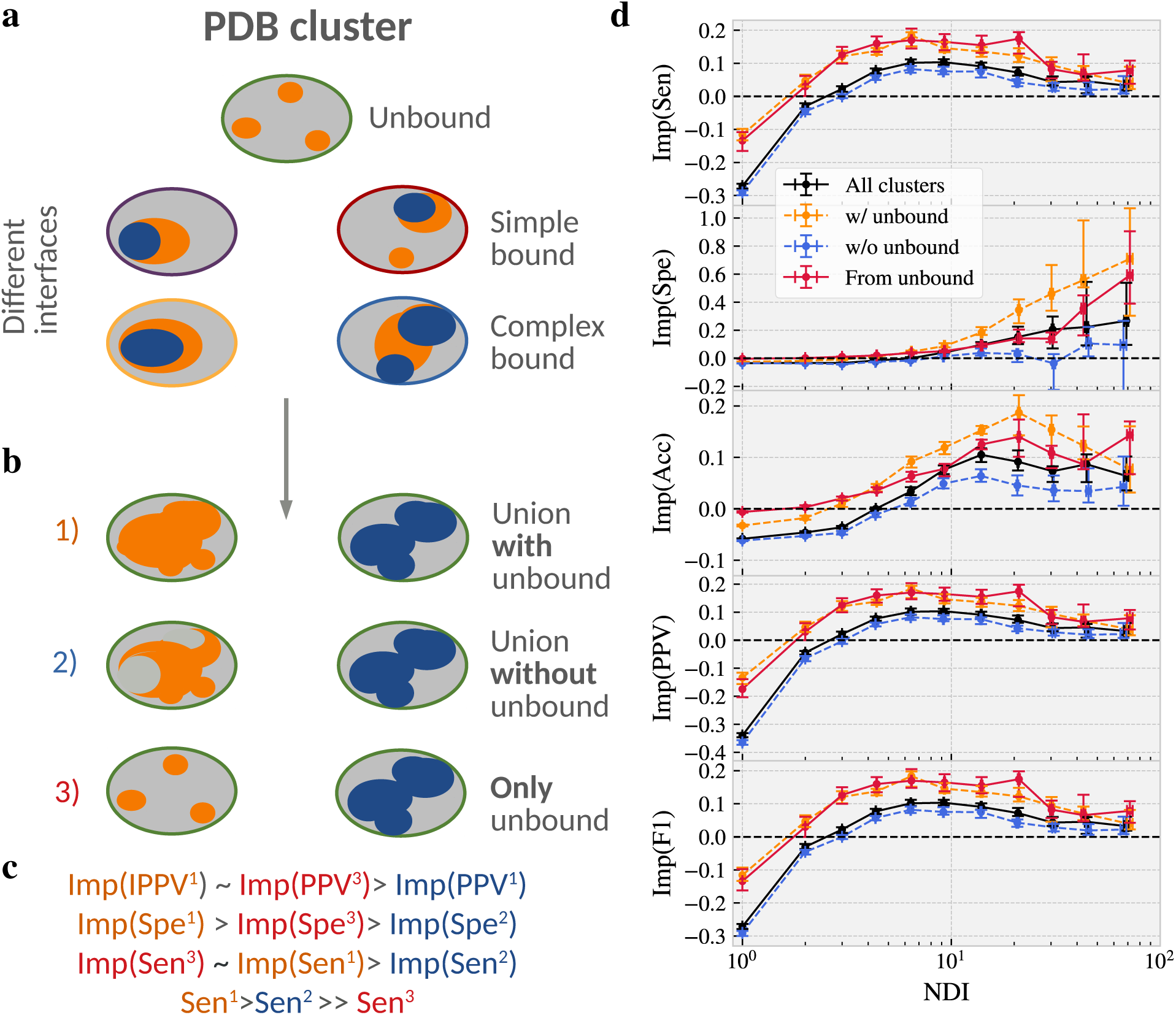
Disorder as a predictor of the assembly path. **a** Sketch of our hypothesis of the role of DRs at the different steps of assembly in a cluster of 4 NDIs. Unbound structures present several disconnected DRs (orange) that mark the location of the possible IRs (blue) accessible from that structure (we call them simple bounds). In the same way, the DRs observed in bound structures, predict the possible IRs of that complex with new partners (what we call complex bound). **b** Sketch of the union of DRs in clusters with, and without unbound structures, and using only the information from unbound structures, compared with the union of all the IRs in the cluster. **c** Expected relative behaviour, under our hypothesis, of the metrics for the groups in b. **d** Median (within bins of similar NDI) of the relative improvement of the predictions based on the disorder with respect to a random guess of the same size for our 5 estimators, see Eq. (1). The median is computed using 4 groups of clusters: (black) all the clusters, as in Fig. 3–**a**, (orange) only clusters with unbound structures, (blue) only clusters without any unbound structure. For the case of clusters with unbound forms, we show the data obtained from predictions based only on the knowledge of the DRs in the unbound (red). The errors give the 90% confidence interval, computed using the bootstrap method with 1000 repetitions.

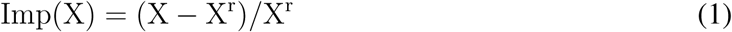

being *X* and *X*^r^, the measured and the randomised values, for our 5 estimators. We show the median of these magnitudes in bins of similar NDI (errors computed with the bootstrap method with a 90% of confidence) in Fig. 4–**b**, and the absolute values in Fig. S4. In all metrics, we get statistically significant improvements of the predictions if the information of the unbound structures is available, notably for the specificity. Yet, the predictions in clusters without known unbound structures are still better than random guesses, but with a particularly poor specificity which is compatible with the fact that we are missing the centre of the interfaces in our predictions since they can only be disordered in unbound forms (see the Sketch of Fig. 4–**a**). The same behaviour is observed for the estimations in clusters with just one interface: if no unbound is present, the predictions are much worse than random because DRs and IRs are generally mutually exclusive in the same structure, an effect much softened if unbound structures are present. It is quite interesting that both the predictions that use the DRs from all the structures or only from the unbound structures show very similar relative improvements in the sensitivity and PPV, even though the predictions of the second group are much smaller and thus worse in absolute value. This tells us that the predictive power of the DRs is, in average, as important in unbound forms as it is in bound ones. All combined, the different relative and absolute behaviours of our 5 estimators in each of our 4 groups of predictions, suggest that DRs change with the interactions (at least at the simplest step of assembly).

Checking our hypothesis at higher steps of protein assembly in a similar statistical way would require, not only homologous families of proteins, but clusters of similar complex structures, which gets much harder at a technical level and will be tackled in future works. Yet, we have tested this idea in particular examples of our dataset. For example, we show in Fig. 5 some of the structures of the cluster 2c8g_A whose UDR and UIR were shown in Fig. 3–**b**. In the centre of Fig. 5–**a**, we show the union of the DRs observed in the two distinct unbound structures of the cluster (2c8b_X and 1uzi_A), and the different simple bound structures observed in the cluster (just one partner, either one protein or a compact complex). We can see that the UDR of the unbound predicts to a good extent the location of the new IRs. However, see Fig. 5–**b**, the DRs observed in the bound structures change drastically their the location. In fact, they appear at the same regions where we observe interfaces of the same complex with other new partners.

**Figure 5:**
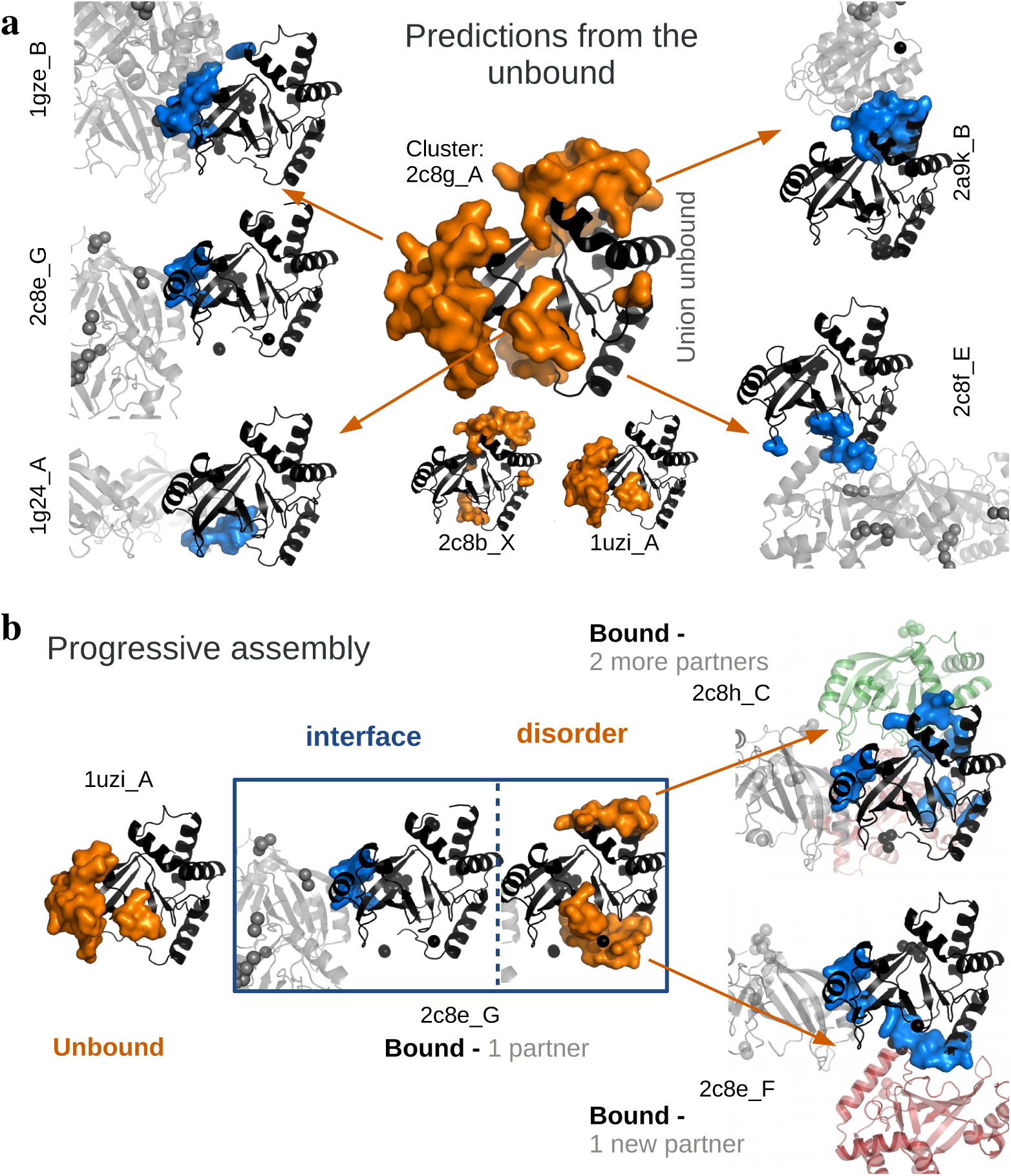
Progressive change of the disorder. **a** We show (in the centre) the union of the DRs of the two distinct unbound structures in the cluster 2c8g_A (in orange), together with the cluster structures with a simple bond (interface region in blue). The structure of the chains that belong to the cluster are shown in black and have all been aligned with the representative’s one (2c8g_A). The structures of the partners are shown in grey. Spheres represent the missing residues, whose position was modelled in the vicinity of the position of its known (sequence) neighbours. **b** We show the progressive change of the DRs upon binding. The DR of one of the unbound structures allows us to predict the interface of one of the simple bound structures (2c8e_G). The DRs observed in this bound structure (square in the centre) predict the additional bindings of this complex with new partners (1gz_C, 2c8h_C, depicted in red and green).

This observation of the change of DRs at different steps of complex formation suggests that disorder might play a role in the order of assembly. To test this idea, we looked for biologically verified orders of assembly of complexes where we could find all the intermediate structures in the PDB. Considering that the disorder depends on the crystallised structure of the complex, it is important to check that the complete (and not a partial, which is the common case) structure of the complex is available. We have manually identified only two different processes where this was possible. The first, taken from Ref.^28^, concerns the verified hierarchical assembly A > AA′ > AA′ A″A‴ > AA′ A″A‴EE′, with *A* and *E* two different protein chains. We show in Fig. 6–**a** the measured DRs and IRs in structures at the 4 different steps of the assembly, showing that IRs appear roughly in the same areas where DRs were observed in structures of the previous step. In Fig. 6–**b**, we show the second example taken from Ref.^29^. According to Ref. ^29^, dimers combine preferentially in two forming a structure with D2 symmetry. We show the DRs in one of the dimers discussed in that paper and compare with the IRs measured in two complexes with that geometry, showing a surprisingly good agreement.

**Figure 6:**
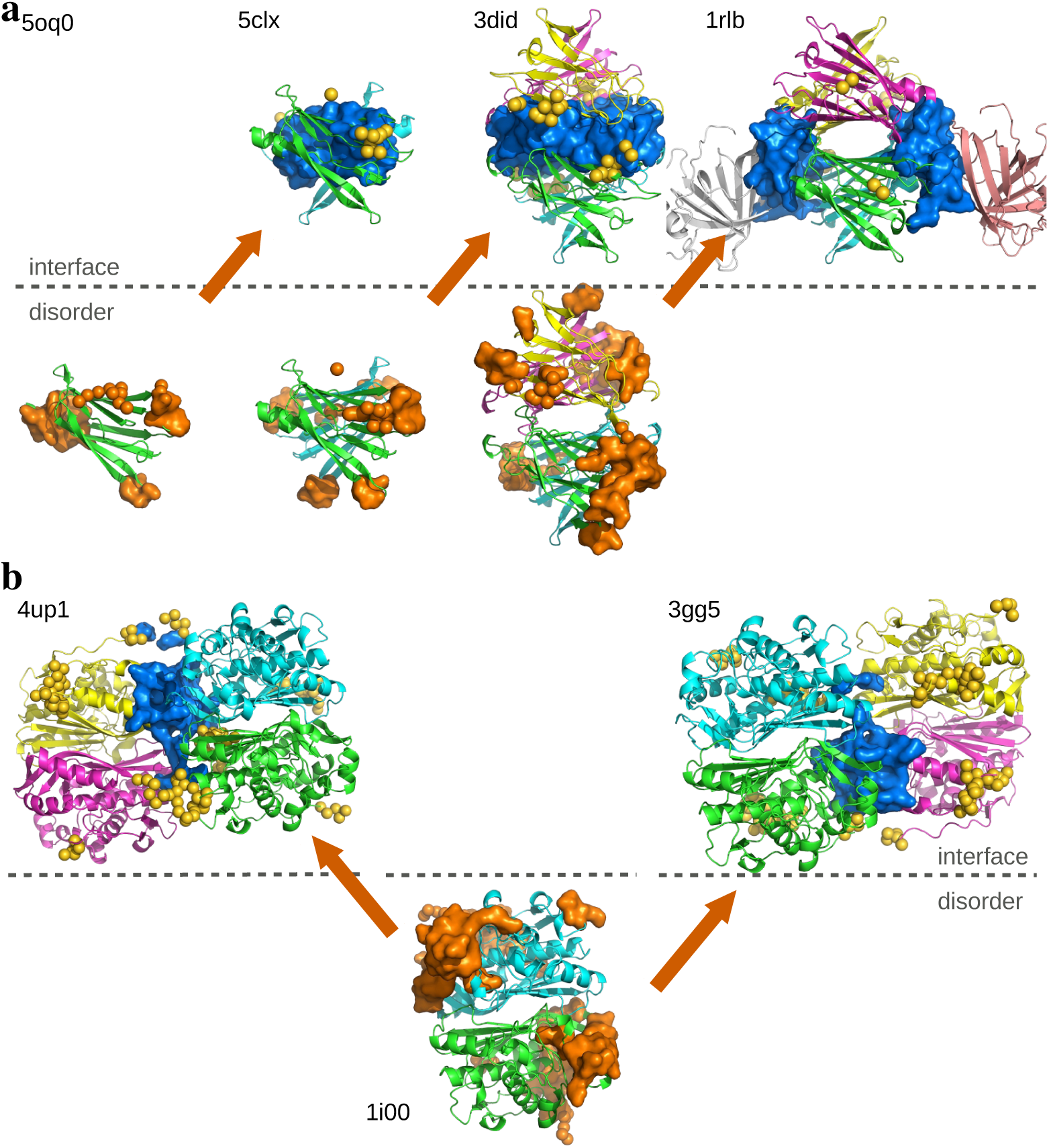
The role of disorder in the biological order of assembly. Evolution of the DRs (orange) and IRs (blue) at different steps of two verified assembly processes. **a.** The assembly of the retinol binding protein complexed with transthyretin (1rlb). **b.** The assembly of the two identical dimers of the human thymidylate synthase (1i00). PDB IDs are reported for each complex.

## 3 Discussion

We have combined all the crystallographic structures available for a large number of homologous families of proteins to conclude that interfaces occur preferentially in regions that are characterised as disordered in alternative crystals. Our analysis uses all the crystallographic structures available in the PDB and does not include any kind of fine-tuning of parameters nor predictions from learning algorithms, which both attaches a great confidence to the generality of the results and leaves great room for improvement. Despite the fact that the database is very incomplete, and thus most of the disorder predictions should seem random in our analysis, we still observe a significant improvement over randomised tests in all the estimators checked. We show that the location of disorder regions changes upon progressive binding. In particular, we show that the knowledge of the disorder of the unbound structures improves significantly the quality of the predictions when compared with clusters without unbound forms. The ratio between the predictor estimators in different groups of clusters allows us to establish a connection between the change in the disorder and progressive binding. We show some examples of how disorder predicts the location of new interfaces from a given structure, which suggests that disorder might have a role in determining the order of assembly of complexes. We verify this idea in the assembly of two complexes whose biological order is known. A statistical study of the change of the disorder in different steps of hierarchical binding is left for the future works.

## Methods

### Database

We include all protein structures in the PDB^26^ (up to the 14/06/2019) available in the old format (which excludes the very large complexes) and with AA sequences of at least 20 residues. We group together all the chain structures with similar AA sequences: a minimum of 90% of sequence identity and length. Clusters are created using the MMseqs2 method^30, 31^, the full list is given in SI section S1. The representative chain of the cluster gives name to the cluster and is chosen as the one coming from the experiment with higher resolution or R-value.

### Interface computation

Protein-protein binding sites are computed using the INTerface Builder method^32^ (two AAs are considered in contact as long as their two *C*_*α*_ are at a distance ≤ 5Å). To obtain the protein-DNA/RNA binding sites, we looked for the residues whose relative surface area (RASA) decreased upon binding. The change in RASA is computed with naccess^33^ (with a probe size of 1.4Å).

### Disorder computation

Our definition of DRs covers all AA whose position is either unknown or poorly known. This means that we include, both the so-called *missing* residues, included in the “REMARK 465” of the PDB structure, and the AA with very high b-factor. We consider a residue has a high b-factor if the experimental b-factor for its *C*_*α*_ atom is beyond one standard-deviation from the mean b-factor of the entire chain. We discuss would happen if only the missing residues were considered in the definition in the SI. In general for all the figures in the paper, and mostly for the sake of a fair comparison between the real and the randomised predictions, we have excluded from the list of sequence AAs, the missing residues that are missing in each and every one of the cluster structures, because, by construction, we cannot say whether they are or not part of an interface. Their relative presence becomes negligible as the NDI increases and when present, they are almost always in direct contact (in the sequence) with the UIR as we show in the SI.

### Union computation

The disordered and binding sites computed at each structure of the cluster are mapped each to the sequence of the cluser representative via sequence alignment using the Biopython’s^34^ PAIRWISE2.ALIGN.GLOBALXX routine. The NDI is the minimal number of such sequences where the number of not shared interface sites is greater than the 5% of the sequence length (compared with all the different interfaces). All the sites that were marked as DR or IR at least once in the cluster are included in the union of DRs and IRs, what we call UDR and UIR respectively.

### Goodness of the predictor estimators

We have considered 5 different estimators, which are computed from combinations of the number of true positives (TP), true negatives (TN), false positives (FP) and false negatives (FN):

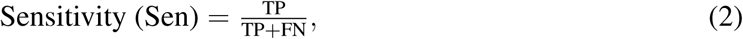

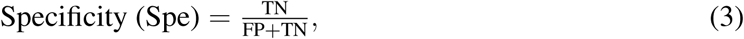

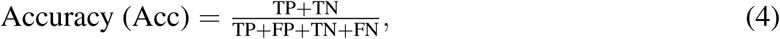

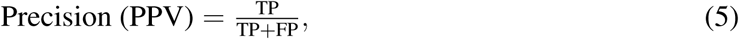

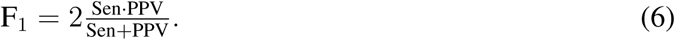

A random prediction of the same exact number of disordered sites, *N*_D_, in a chain of *L* AAs, would predict correctly an interface site (in average) with a probability of *r*_I_ = *N*_I_*/L*, being *N*_I_ the number of interface sites in the cluster. This means that one expects, in average,

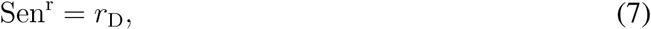

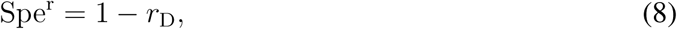

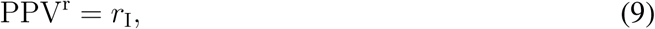

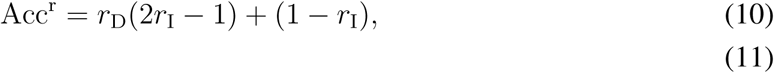

being *r*_D_ = *N*_D_*/L*.

## Data availability

All data necessary to reproduce the analysis (list of clusters of homologous chains, list of different interfaces, list of clusters with unbound structures, list of properties for each cluster) can be found online at www.lcqb.upmc.fr/disorder-interfaces/.

## Acknowledgements

We thank Flavia Corsi and Elodie Laine for useful discussions and technical assistance during early stages of this project. This work was supported by LabEx CALSIMLAB (public grant ANR-11-LABX-0037-01 constituting a part of the “Investissements d’Avenir” program – reference : ANR-11-IDEX-0004-02); BS was partially supported by Ministerio de Economía, Industria y Competitividad (MINECO) (Spain) through Grants No. FIS2015-65078-C2-1-P and PGC2018-094684-B-C21 (also partly funded by the EU through the FEDER program).

## Author contributions

B. S. and A. C. conceived and designed the study and wrote the manuscript. B. S. performed the data analysis.

## Competing Interests

The authors declare that they have no competing financial interests.

## Supplementary information

is available for this paper at REF.

## Notes

https://www.lcqb.upmc.fr/disorder-interfaces/

