## Supplementary Information for "Disorder and interfaces in proteins are two sides of the same coin"

**Beatriz Seoane**
Sorbonne Université, CNRS, IBPS,
LCQB - UMR 7238, and ISCD,
4 place Jussieu, 75005 Paris, France.

**Alessandra Carbone**
Sorbonne Université, CNRS, IBPS,
LCQB - UMR 7238,
4 place Jussieu, 75005 Paris, France.

### S1 About the clusters composition

The list of clusters used, together with the PDB index of the structures of the chains included, the list of structures with the different interfaces in each cluster and the list of clusters with unbound forms can be downloaded online `www.lcqb.upmc.fr/disorder-interfaces/`.

A breakdown of the number of clusters considered in each group of clusters discussed in the paper is shown in Table S1, where we can see the amount of clusters that contain at least one interface and the number of PDB complexes and individual chains. In Fig. S1, we include some additional histograms concerning the number of different interfaces and unbound structures.

### S2 List of structures of cluster 6c9c_A (Fig. 1 main-text)

We show in Fig. 1 of the main text (MT) 13 structures for the protein chains in the cluster 6c9c_A. One of them is an unbound structure (chain A of PDB index 6c9c), and the other 12 cover are the different interfaces measured in the cluster. Their PDB indexes and chain names are 5n8c_A, 5n8c_B, 4fw3_A, 4fw3_C, 4fw4_A, 4fw4_C, 4fw5_C, 4fw7_A, 2ves_A, 3u1y_A, 3u1y_B, 4okg_A.

### S3 Total randomisation test of the disordered regions

In this section we compare the statistics and prediction power of the disordered regions (DRs) and with the random guess. With this aim we proceed as follows. We read the disordered (as missing residue or high b-factor) aminoacids from each chain structure in the cluster and randomise its location in the sequence. We consider two possible of such randomisations: a purely random one but not very meaningful in a biological sense, namely **(test 1)**: a random reshuffling of the DRs (by means of a random permutation of all the sites in the sequence). Mind that such a reshuffling losses one of the main characteristics of the interfaces, the existence of compact regions. We also test a more biologically motivated one, namely **(test 2):**, We keep fixed at their position the disordered aminoacids at the N-term and C-term (which are known to be widely present all along the PDB), and we randomly displace internal disordered sites, keeping sequential disordered sites always together. After these two randomisations, we compute the DR as the union of all these random disordered sites in the cluster, as done in the MT, and compare its properties with those of the experimental IRs and its power to predict IRs. In Fig. S2, we show the mean relative size of these DRs for the two sets. Clearly, the behaviour of these fake disordered regions with the number of interfaces in the cluster, whose sizes cover essentially the

| | Clusters | Clusters w/ interfaces | Complexes | Chains |
|---|---|---|---|---|
| All clusters | 47102 | 36628 | 134337 | 354420 |
| Clusters w/ unbound | 16605 | 6131 | 82143 | 133526 |
| Clusters w/o unbound | 30497 | 30497 | 58042 | 220894 |

Table S1: Breakdown of the number of clusters.

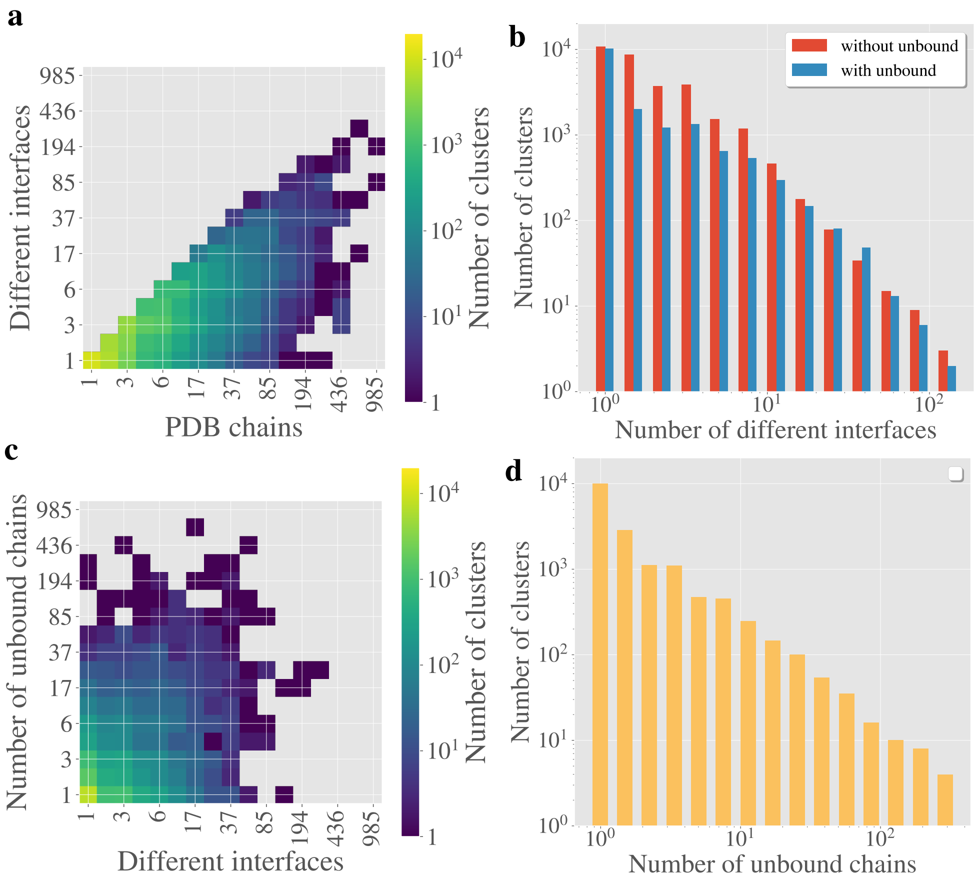


Figure S1: **a** Bi-dimensional histogram of the number of clusters with a given number of different interfaces and a given size. **b** Number of clusters as function of the number of different interfaces for the groups of clusters that either contain any unbound chain or they do not. **c** Bi-dimensional histogram of the number of unbound chains in the cluster as function of the number of different interfaces. **d** Number of clusters as function of the number of unbound chains contained in each cluster (only clusters with unbound structures considered).

whole chain above 10 interfaces, differs strongly with the data shown for the union of the interface regions (or the real disordered regions).

### S4 Number of connected regions in randomised tests

We compare the data shown in Fig. 2 in the MT, concerning the number of connected interface or disordered regions of our clusters (normalised by the sequence size), with the number of clusters that one would measure if the DRs were random. We consider two distinct tests: (**test 3**), a random permutation of the DR sites in the sequence, and (**test 4**), a reshuffling of the DRs keeping together all sequential disordered sites, keeping in both cases the total number of disordered sites fixed. We show in Fig.S3 the averaged number of this number of connected regions among all the clusters with the same number of different interfaces. Again, both tests give significantly different curves than the real ones, as long as the NDI is not too high (where the disordered regions superimpose forming one or two very large clusters).

### S5 Absolute values of the estimators for the unbound/not unbound groups.

We show in Fig. S4 the same data shown in Fig. 3 in the MT but this time we plot the absolute value for the estimators instead of the improvement with respect to a random reshuffling.

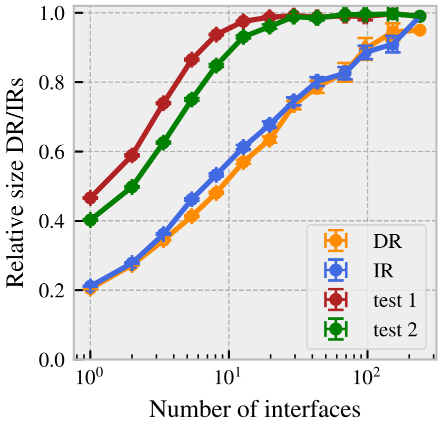


Figure S2: **Total randomisation test.** We compare the averaged relative size of union of disordered (orange) and interface (blue) regions with respect to the sequence length as function of the cluster size, with the averaged relative size of the union of fake disordered regions obtained after reshuffling the experimental disordered regions (test 1, red) and after reshuffling only the internal disordered domains and keeping sequential groups of amino-acids together (test 2, green). In both tests, the randomised disordered regions follow a rather different behaviour with the number of interfaces than the union of experimental interface regions.

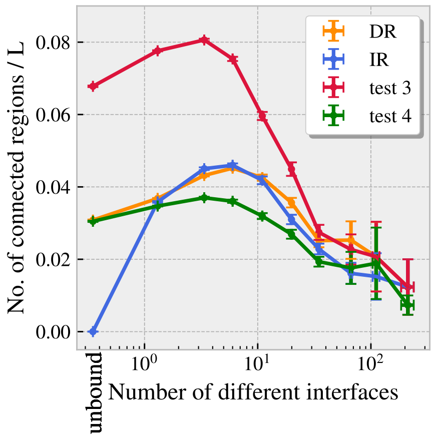


Figure S3: **Number of connected regions in reshuffling tests.** We compare the averaged number of connected disordered regions (DR, orange; Fig. 2b) and interface regions (IR, blue; Fig. 2b), normalised by the sequence length, as function of the number of different interfaces in the cluster with the numbers we would obtain if the same number of disordered sites where randomly distributed. We have considered two distinct randomisation tests: a random permutation of the disordered sites in the sequence (test 3) and a reshuffling of the disordered regions but keeping consecutive disordered sites together. Both tests lead to different curves than the real ones, with the exception of the very big clusters, where the regions superimpose forming a very large cluster.

### S6 Test F1 compared to other reshuffling tests.

We discussed in the MT, that our estimators of the goodness of the prediction of the UIR based on the knowledge of the UDR of each cluster should be compared with the values one would obtain if the disordered residues in the UDR were randomly distributed in the chain (but with the total number of them missing). In particular, in the MT, we made comparisons of the real data measured in the cluster, with the averaged values one would expect to obtain, cluster-by-cluster, if these residues were completely random (see Methods and test 3 of Fig. S3). Here we compare our results with other kinds of randomisations one could think of, in particular, randomisations that would preserve the quality of having compact DR regions (just as we observe for the IRs and the real DRs). We considered two methods

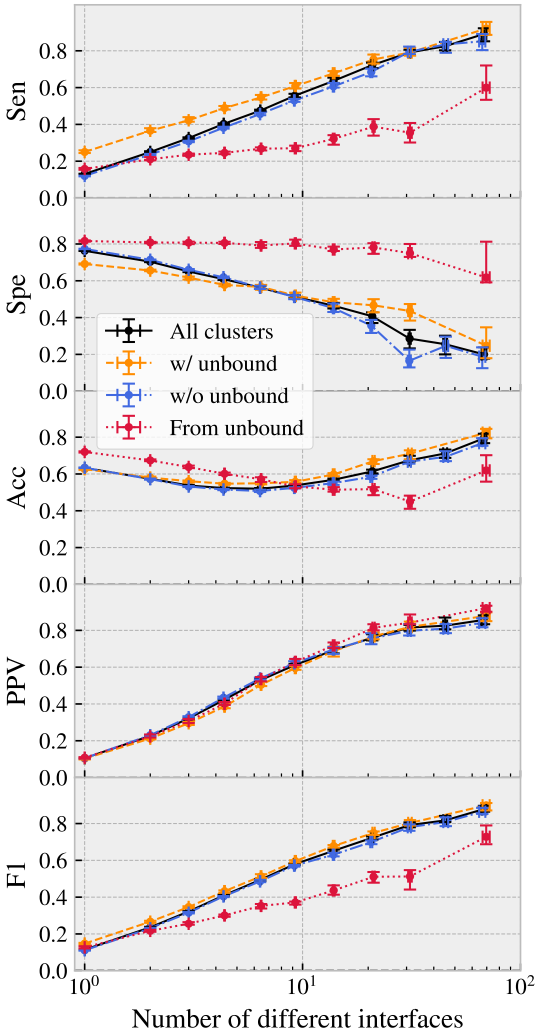


Figure S4: **Effect of the unbound forms on the goodness of the use of disorder as a predictor for interfaces.** We show the absolute values of the estimators shown in Fig. 4-**b** in the MT for the same predictions, that is, using either the total union of disorder regions averaged in all clusters (black), only clusters with unbound forms (orange) or only clusters without unbound forms (blue). We also consider the predictions of the union of disorder measured only in the unbound forms (red).

to do this: (**test 4**, as in Fig. S3) we reshuffle together the groups of sequential disordered residues, or **(test 5)** we draw randomly in the sequence groups of sequential disordered residues, with random length each. In both cases, it is important to ensure that the total number of disordered residues after the reshuffling is preserved. We show the difference between the real value and the randomised version of the 5 estimators for all these three tests in Fig. S5 (for the groups of clusters with unbound forms), showing very similar results.

### S7 Disorder regions only as missing regions

In this section we discuss what would be the result of the analysis if we restricted ourselves to identify as disordered only the missing residues. We proceed as in the MT: we identify the missing residues at each structure of the cluster, map it in the cluster's representative sequence and compute the union of all these regions that were at least once identified as missing in the cluster. We show an example of this procedure in the cluster 6c9c_A in Fig. S6–**b**. We call this, union of missing regions (UMR). Like in Fig. 2 of the MT, we show in Fig. S6–**a** the distribution of the relative sizes of the UMRs, $r_{\mathrm{M}}$ as function of the NDI, in yellow, and compare it with the relative size of the UIRs, in blue. As for UDRs,

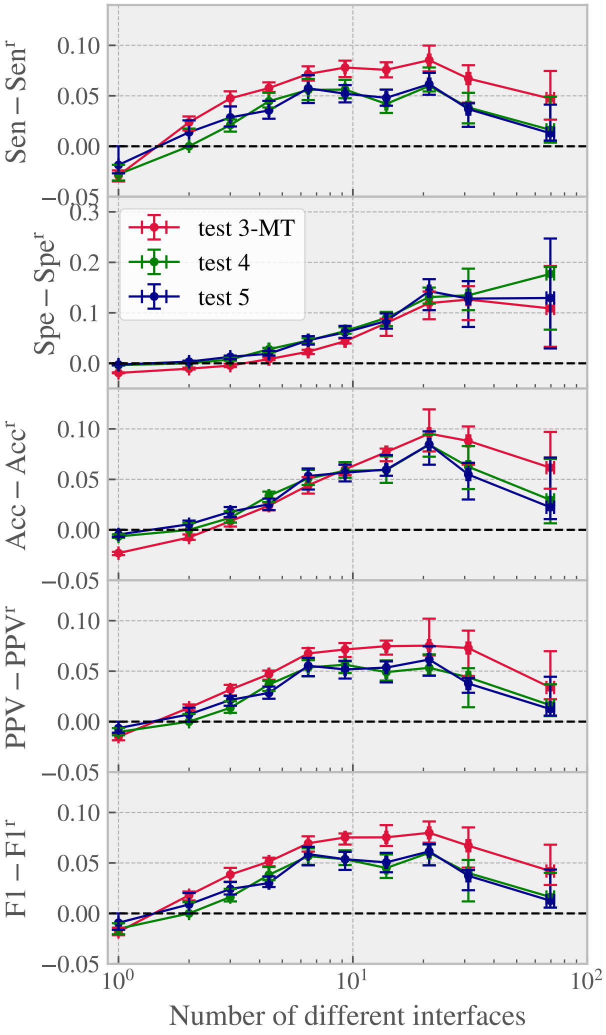


Figure S5: **Improvement of the estimators' measures with respect to different reshuffling tests**. We show the median of the difference between the real and the randomised (see text for a description of each test) values for the estimators, cluster-by-cluster, and computed within groups of clusters of similar number of interfaces. For the sake of clarity, we just considered the clusters that have both unbound and bound forms. Errors are computed using the bootstrap method with 1000 repetitions and a 90% of confidence.

$r_{\mathrm{M}}$ grows with the NDI at a similar rate than $r_{\mathrm{I}}$, but its total size is much smaller than the other two. In Fig. S6–**c**, we show, for each cluster, the value we obtain for our 5 estimators of the quality of a predictor based on the missing residues. As in Fig. 3-**a** of the MT, we compute the median of these values using bins of clusters of similar NDI and compared this curve with the one we would expect if the UMR was randomly distributed along the sequence. As we can see, the PPV is much higher in the real than in the random case. Yet, the UMR covers very small portions of the interfaces, which leads to very poor gains for the rest of the estimators.

### S8 Fraction of forever missing residues

In general, for the analysis, we have removed from the total list of AA of each cluster, the missing residues that are missing in all the structures of the cluster, because we cannot determine if they belong or not to an interface. The portion of this forever missing residues becomes extremely small as NDI grows, see Fig. S7.

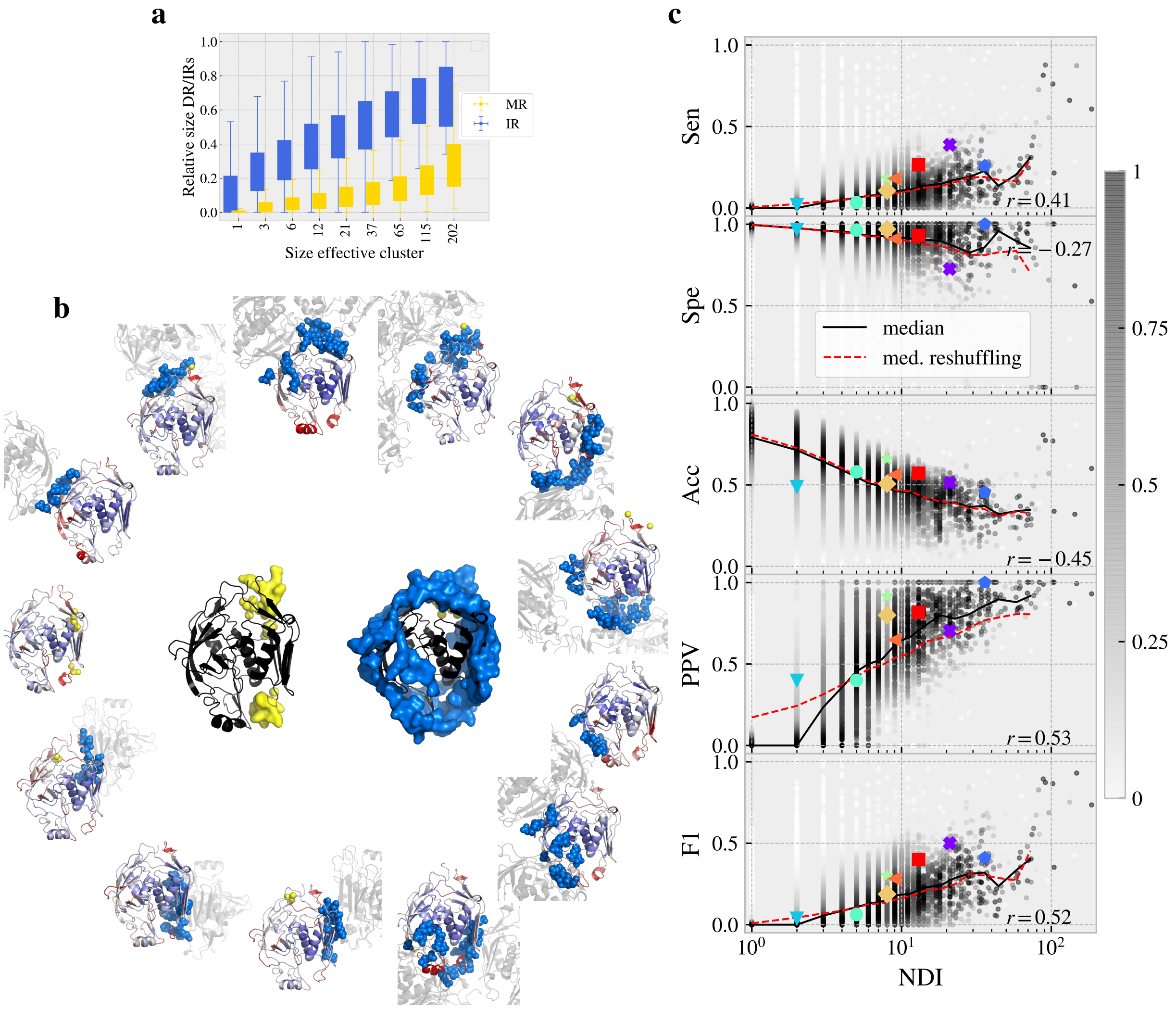


Figure S6: **Union of missing regions**. **a** We show the relative size of the union of the missing regions as function of the NDI. **b** In the external wheel, we shown an example of the structures that compose the cluster 6c9c_A with their missing and binding sites marked as yellow and blue spheres, respectively. The structure is coloured with the normalised b-factor, being red and blue, high and low b-factor, respectively. Partners are shown in light grey. In the centre, we show the union of missing residues (yellow surface) and the union of interface regions (blue surface). Compare to Fig. 1. **c** The measured value of our 5 (goodness of the predictor) estimators for all clusters available in the PDB. The median of these values is shown as a black line and the median of the values one would obtain if the UMR sites were randomly distributed is shown in red. Colored big dots refer to those complexes studied in Figs. 3–a and 3–b.

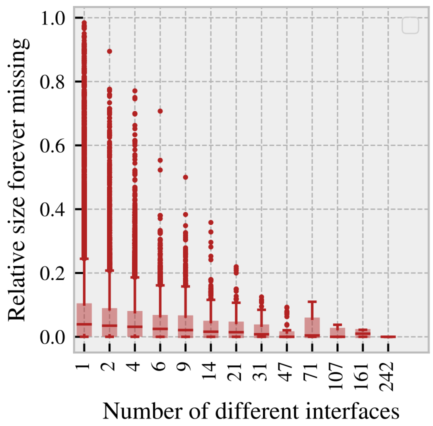


Figure S7: **Forever missing residues.** We show a box-plot of the fraction of residues in the cluster sequence that is reported as missing in all the structures of the cluster as function of the NDI. Data was analysed in bins of similar NDI (bins distributed evenly in the logarithmic scale).
